# Utilizing blood single-cell transcriptomics to integrate intrinsic and systemic immune aging

**DOI:** 10.1101/2024.10.30.621198

**Authors:** Alan Tomusiak, Sierra Lore, Morten Scheibye-Knudsen, Eric Verdin

## Abstract

Biomarkers of aging provide insight into the biological effects of interventions and diseases. However, most biomarkers today are based on measurements derived from bulk cell measurements, making it challenging to interpret whether an effect is due to changes in cell type composition (systemic factors) or a cell intrinsic effect. Single-cell RNA sequencing provides a unique platform to simultaneously compare aging-associated changes on both a cellular and bulk level. We first generated a single-cell combined automated human blood cell type and age predictor (clock) for six distinct human T cell subsets. We applied these tools to find acute COVID is associated with a shift in CD8+ cytotoxic cell proportions, while cell type proportions are stable in patients with HIV on long-term ART treatment (HIV+ART). Both COVID and HIV+ART were associated with an increase in naive CD8 T cell transcriptomic age. We further found our single-cell aging biomarker is linked to ribosomal gene expression and has a link to mean cellular transcript length. This study highlights the potential of single cell transcriptomic biomarkers for understanding how the human immune system is impacted by age-associated systemic changes in cell type composition and intrinsic cellular aging.

## Introduction

Aging is a complex, systemic process that involves changes in cellular function, cell composition, tissue organization, and intercellular communication networks^1–3^. Developing biomarkers capable of measuring aging on each level of organization is critical for understanding how molecular and physiological changes occur across the human lifespan. Although numerous tools have been designed to quantify aging, most existing biomarkers focus on a single level of organization. This limitation hampers our ability to grasp the intricate interplay between cell-intrinsic and systemic aging. Creating novel interpretable tools that capture this complexity is critical for the development of interventions to improve human healthspan and lifespan.

Early efforts to design accurate aging biomarkers, often referred to as “clocks,” focused on single molecular measurements. The first clocks were identified by Hannum and Horvath and colleagues based on changes in DNA methylation in 2013^4,5^. These clocks predicted biological age using CpG markers derived from DNA methylation data. Over time, clocks were developed based on bulk transcriptomics^6^, proteomics^7,8^, ATAC-Seq^9^, and other molecular measurements. These advancements have led to new methods for understanding aging across different levels of resolution, each providing unique insights into the aging process. However, the interpretation of these biomarkers can be challenging due to their reliance on measurements derived from a bulk collection of cells or plasma.

To solve this challenge, numerous groups have begun to develop clocks based on single-cell measurements. In 2021, Trapp et al. developed a clock based on single cell methylation data^10^. In 2022, Buckley et al. used single-cell transcriptomic profiling data to predict chronological and biological age^11^. Using transcriptomics instead of DNA methylation identifies markers that are easier to link to changes in specific proteins and pathways thereby providing novel insights into changes that occur with biological age as well as allowing for analysis of new datasets^12,13^. Furthermore, cell-type specific effects of aging and rejuvenation can be measured. As an example, Yu et al. (2023) used a technical innovation on single-cell transcriptomic data to derive novel insights regarding the effect of exercise on neuronal rejuvenation^14^. Similarly, single-cell chromatin measurements based on imaging have recently been developed to quantify age reversal by partial reprogramming^15^. This development enables scientists to explore how individual cells within an organism age differently, contributing to our understanding of cellular heterogeneity in aging.

Concurrently, several groups developed biomarkers to predict aging using systemic measurements. Initial efforts were focused on predicting systemic measurements using molecular readouts. As an example, Levine et al. (2018) used methylation data to predict composite clinical data to generate a clock capable of predicting several key healthspan measurements. GrimAge further improved upon this by predicting time to death, time to coronary heart disease, and time to cancer using a similar technique^16,17^. More recently, clocks have been developed to predict the aging of individual organs^18^ and physiological systems^19^. These measurements allow for better understanding of systemic changes directly relevant to human health.

Several studies have integrated aging on several levels of resolution simultaneously, frequently by employing multi-omics. In the context of human aging, these measurements have led to improved clinical predictions^20^ and the discovery of age-associated chemokines^21^. Other groups have combined multi-omics with longitudinal data, enabling a deeper understanding of the dynamics of human aging^2,22^. Combining measurements of systemic aging with advancements in single-cell biomarkers has the potential for unlocking a deeper understanding of the interplay between extrinsic and intrinsic aging.

Simultaneous profiling of intrinsic and systemic aging has particular importance in the context of the immune system^23^. Due to both intrinsic (i.e. cell autonomous) and extrinsic factors (for example, thymic involution), the naïve CD8+ and CD4+ T cell compartments decline over time^24^. These can interfere with the body’s ability to repair tissues and resolve novel infections^25^. In addition to the decline in intrinsic cell function, the immune system also experiences prominent age-related changes in cell-type composition – for example, via a decline in naïve CD4 and CD8 T cells and a concomitant increase in memory and effector T cells^23,26–28^. This complex interplay of cellular and systemic aging is important, but our ability to assess how each of these independently contributes to aging-associated pathology is presently limited.

Here, we use a previously published single-cell transcriptomic dataset^23^ to generate predictors of T cell composition and individual cellular aging. We demonstrate that our cell type predictor can identify and quantify six canonical T cell subsets (naïve CD8s, central memory CD8s, effector memory CD8s, naïve CD4s, central memory CD4s, and regulatory T cells) and their changes in relative abundance during aging. Using this cell type predictor, we then generated six individual age predictors for each predicted T cell subtype. We applied our cell type and aging predictors to two datasets – one of acute COVID and another of HIV-infected individuals with long-term ART treatment. Similarly to what we have shown previously using epigenetic data^29^, we find that acute COVID is associated with changes in cell type composition. We further show that both diseases are associated with intrinsic accelerated aging in naïve CD8 T cells. Lastly, we investigate the mechanistic drivers of our age predictors and identify associations with ribosomal gene expression and mean cell transcript length.

## Results

### Automated prediction of cell type recapitulates known changes in T cell composition with age

To simultaneously profile changes in blood cell type composition and intrinsic age-associated changes, we created separate models: a cell type prediction model and six age prediction models for each cell type (naïve CD8s, central memory CD8s, effector memory CD8s, naïve CD4s, central memory CD4s, and regulatory T cells). Automating cell type prediction allows for less bias during cellular age prediction. As the basis for our model, we used a previously published scRNA-Seq dataset of two million peripheral blood mononuclear cells (PBMCs) from 166 individuals^23^. As noise can significantly affect accurate cell type or age prediction, we filtered our analysis on genes showing some correlation (R > .01 or R < -.01) with age. This removed approximately 90% of genes in the original dataset (Figure 1a).

**Figure 1.**
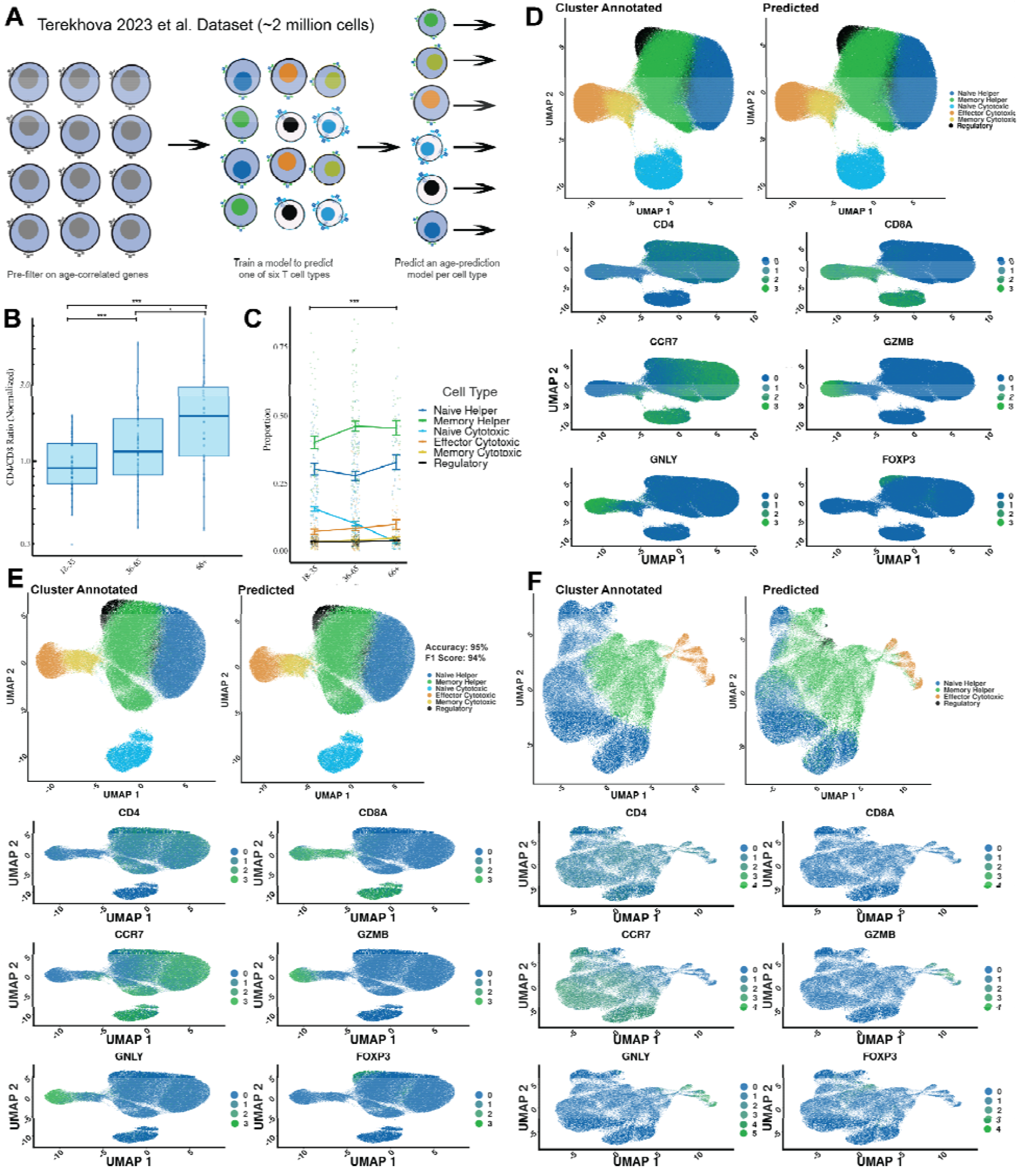
Cell type predictions recapitulate known effects of aging on the immune system. **a**, General study and model design. **b**, CD4/CD8 ratio identified in different age groups, normalized for ratio identified in individuals between 18-35. **c**, Cell type proportion changes in individuals from different age groups. **d**, Comparison between predicted and manually annotated cell clusters for the training set. CD4, CD8A, CCR7, GZMB, GNLY, FOXP3 are used as ground truth markers for cell type labeling. **e**, Comparison between predicted and manually annotated cell clusters for the test set. **f**, Comparison between predicted and manually annotated cell clusters for an external validation set (Yasumizu et al., 2024). *** Bonferroni-corrected p-value less than or equal to .001, * Bonferroni-corrected P-value less than or equal to .05, # Bonferroni-corrected P-value less than or equal to .1.

We then split the dataset on a per-donor basis into a training subset (80%) used for building the model and a test subset (20%) for determining its accuracy and precision. We used an additional external dataset^30^ to measure the ability of our models to make accurate predictions in other cohorts. To determine cell prediction accuracy and generate initial labels, we used K-means clustering to split the cells into biologically relevant groups and then labeled these groups based on the expression levels of six cell surface markers (CD4, CD8A, CCR7, GZMB, GNLY, and FOXP3). Cell subsets were identified based on the following positive markers: naïve CD4 helper T cells (CD4, CCR7), central memory CD4 helper T cells (CD4), CD8 naive cytotoxic T cells (CD8A, CCR7), CD8 effector cytotoxic T cells (CD8A, GZMB, GNLY), CD8 memory cytotoxic T cells (CD8A), and regulatory T cells (FOXP3, CD4).

We validated our model by first identifying whether previously reported aging-associated trends could be recapitulated. In agreement with previously identified and reported changes^28^, we identified an increase in the CD4/CD8 ratio with age (p=.0003) (Figure 1b). Within the six cell types we measured, we observed a significant decrease (p < .0001) in the proportion of CD8 naïve cytotoxic cells (Figure 1c) with age, which is also in accordance with previous literature^31^.

We further tested our cell type prediction model by comparing predicted cell types to those we manually annotated. In both the training (Figure 1d) (97% accuracy; .98 F1 score) and test (Figure 1e) (97% accuracy; .97 F1 score) datasets, the predicted cell types closely match the manually annotated clusters. Furthermore, there is accordance with cell types as identified by canonical cell markers CD8A, CD4, CCR7, GZMB, GNLY, and FOXP3. We then tested our cell type prediction model using an external dataset of CD4+ T cells^30^. We identified strong accordance (83% accuracy; .80 F1 score) between the manual annotation and assumed clusters based on canonical cell markers (Figure 1f).

### Cell type-dependent models predict age across a variety of cell types

After validating our cell type prediction model, we then trained six independent age prediction models, one per each of the six predicted cell types (naïve CD8s, central memory CD8s, effector memory CD8s, naïve CD4s, central memory CD4s, and regulatory T cells). We then compared the predicted age to the chronological age of the donor. We both compared the predicted age of the individual cell to the donor it derived from, as well as the mean average predicted cell age per donor.

For the training set, we observed a strong correlation between the predicted age of a cell and chronological age (R= .56, mean absolute error (MAE) = 11.9, p < 1*10^−16^), particularly when the average age of all cells per donor was calculated (R= .84, MAE = 11.4, p < 1*10^−16^) (Figure 2a). Similarly, we observed a moderate-to-strong correlation in the test set (R=.49, MAE = 13.5, p < 1*10^−16^), which also increased when computed as the average age of all cells per donor (R= .8, MAE = 11.5, p < 1*10^−12^) (Figure 2b). We applied our model to the Yasumizu et al. (2024) dataset and observed a weaker but still significant correlation on a cell level (R= .38, MAE = 15.2, p < 1*10^−16^), and when cells were averaged per each donor (R= .78, MAE = 6.5, p = .001) (Figure 2c).

**Figure 2.**
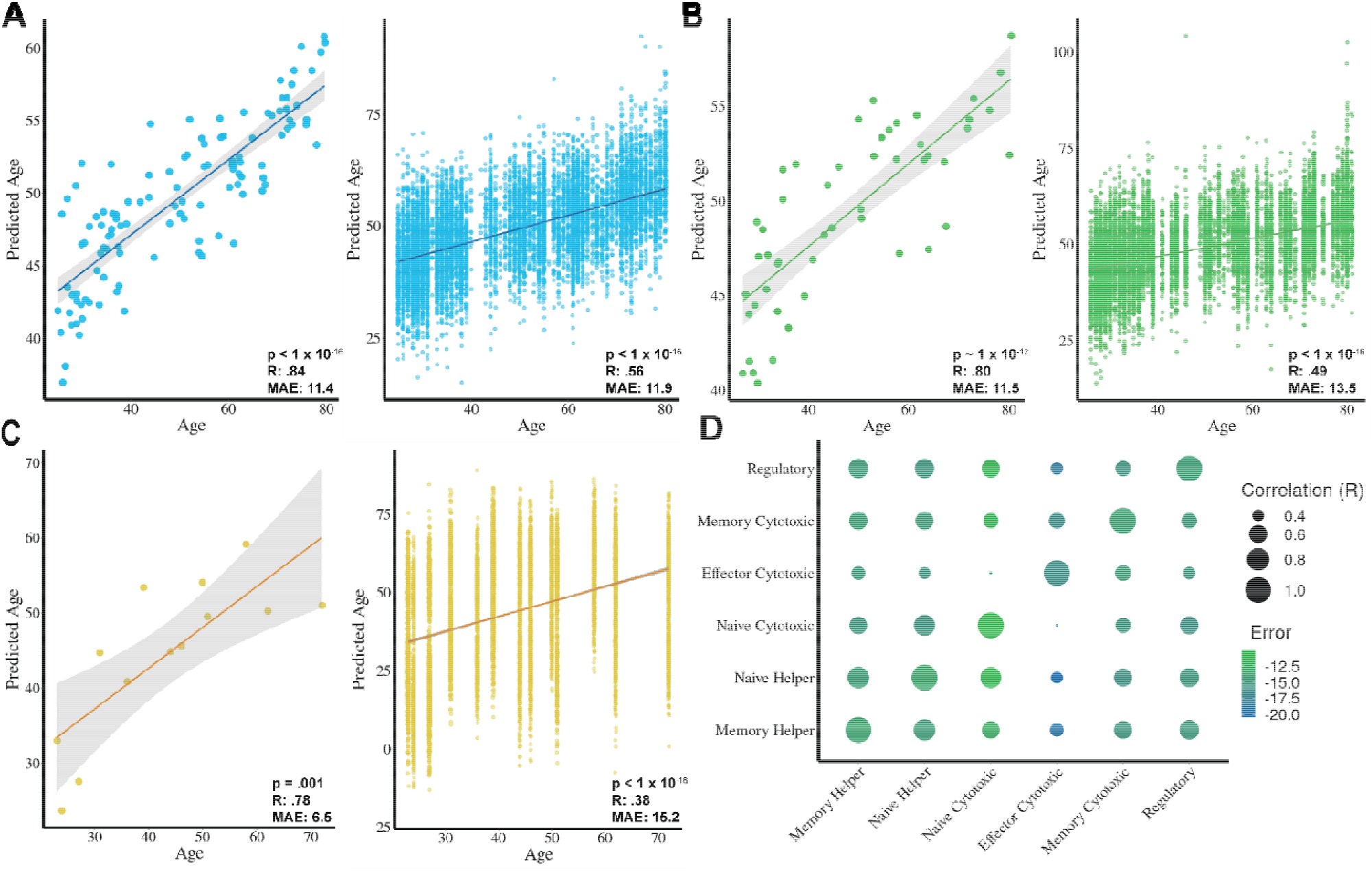
Age prediction of T cells in the training, test, and validation sets. **a**, Predicted age compared to chronological age of donors (left) and cells (right) in the training set. **b**, Predicted age compared to chronological age of donors (left) and cells (right) in the test set. **c**, Predicted age compared to chronological age of donors in the validation set (28). **d**, Correlations of age prediction between each T cell subset clock, and relative errors of each T cell subset clock relative to donor chronological age.

To determine whether each of these age predictors was identifying unique cell type-specific aging patterns, we tested whether they were correlated with each other. In general, the individual age predictors had moderate (R ∼ .5) age prediction correlations with one another, suggesting they measure both a shared and cell-type dependent aging signature (Figure 2d).

### COVID-19 impacts immune aging through both cell composition and intrinsic aging mechanisms

Aging and diseases such as COVID-19 have proven to have significant impacts on T cell composition and immune function^32^. We were interested to find whether acute COVID-19 would impact cell type composition, cellular aging, or both. We used an external COVID-19 dataset made up of 171 COVID-19 infected and 25 healthy individuals^32^, filtering on individuals who had COVID-19 within two weeks of sample collection.

We initially applied our cell type prediction model to this dataset (Figure 3a) and found identification of distinct T cell subsets to be in accordance with cell markers (Figure 3b). Next, using the age prediction models, we evaluated the predicted ages of individual cells and the predicted ages of donors. We found that our models achieved moderate accuracies on individual cells (R= .26, MAE = 16, p < 1*10^−16^) (Figure 3c) and cell ages averaged per donor (R = 0.66, MAE = 14, p < 1*10^−10^) (Figure 3d).

**Figure 3.**
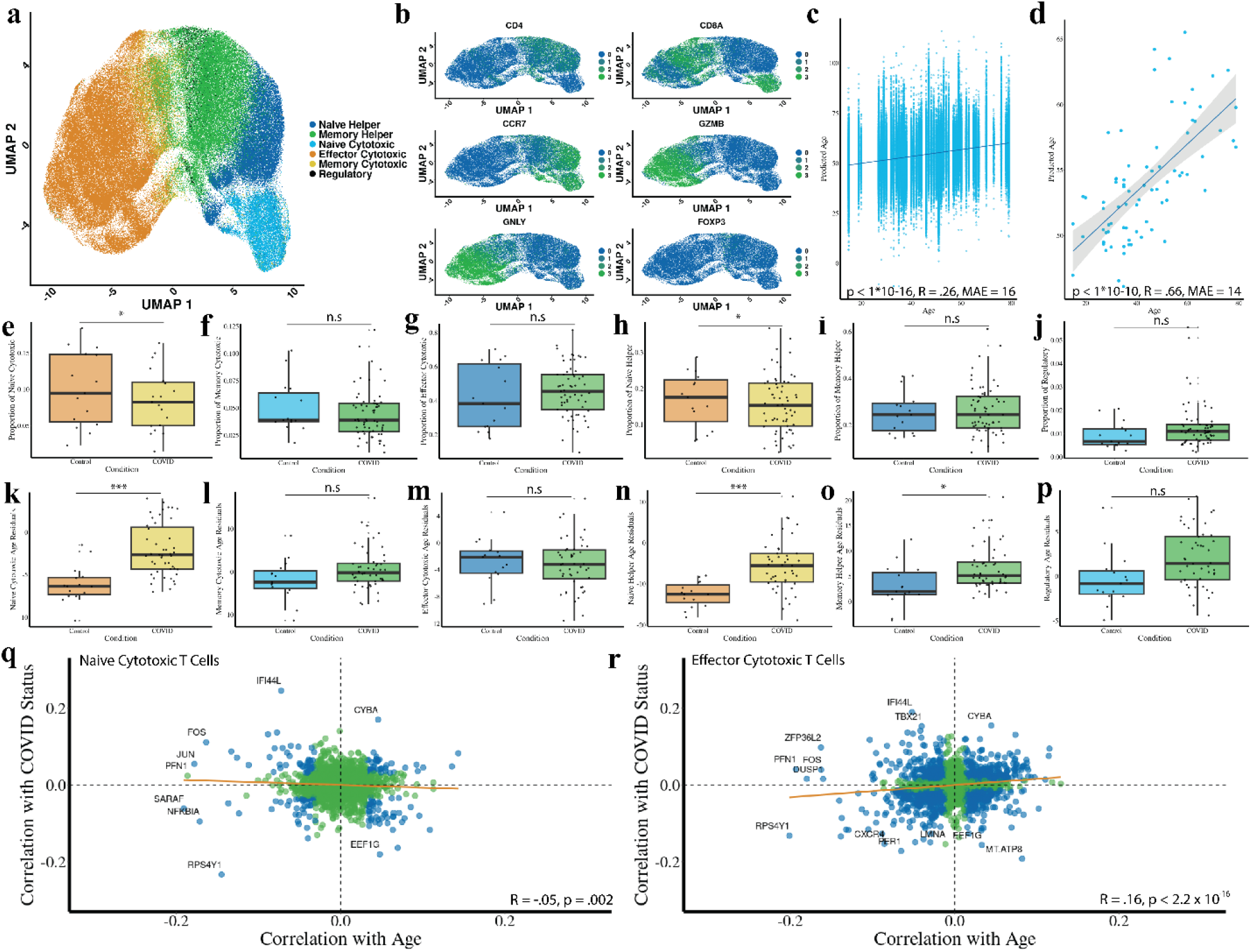
Impact of age and COVID-19 on T cell composition, cellular aging, and gene expression. **a**, Predicted cell types using the cell type prediction model. **b**, Expression levels of canonical cell markers. **c**, Predicted age of individual cells versus donor chronological age. **d**, Mean prediction donor age across all cells compared to donor chronological age. **e-j**, Proportion of cells from COVID-19 donors and from healthy donors by cell type. **k-p**, Residual age prediction by condition for each cell type. **q**, Plot comparing genes correlated with COVID and aging in naïve cytotoxic T cells. Blue represents a statistically significant effect. Orange line represents the mean regression. **r**, Plot comparing genes correlated with COVID and aging in effector cytotoxic T cells. Blue represents a statistically significant effect. Orange line represents the mean regression. *** Bonferroni-corrected p-value less than or equal to .001, * Bonferroni-corrected P-value less than or equal to .05, # Bonferroni-corrected P-value less than or equal to .1.

We then sought to determine whether acute COVID-19 infection affects cell type composition. We observed significant decreases in predicted proportions of naïve CD8 (Figure 3e; p = .03) and naïve CD4 cells (Figure 3h; p =.03). We observed weaker changes in cell type proportions for other cell types, including CD8 central memory cells (Figure 3f, p = .21), CD8 effector cells (Figure 3g, p = .3), CD4 central memory cells (Figure 3i, p = .83), and regulatory T cells (Figure 3j, p= .08). As naïve cells make up approximately 30% of cells in the patient samples, lowering their relative proportion is a significant immunological impact.

Next, we were interested in whether predicted cellular ages are affected by COVID-19. We observed significant age acceleration in naïve CD8 (Figure 3k, p = 1e^-6^), naïve CD4 (Figure 3n, p = 1e^-7^), and CD4 memory (Figure 3o, p=.02) cells. We did not see a significant shift in predicted age for CD8 memory (Figure 3l, p = .2), CD8 effector (Figure 3m, p=.98), or regulatory (Figure 3p, p = .07) T cells. Lastly, we were interested in determining whether there would be a shared transcriptional signature between aging and COVID-19 in naïve CD8 and effector CD8 cells. There was only a slight negative (Figure 3q, R = -.05, p = .002) and a slight positive (Figure 3r, R = .16, p < 1*10^−16^) correlation between COVID and aging in naïve CD8 and effector CD8 cells, respectively.

### Long-lasting ART treatment and HIV impacts aging through cell-intrinsic mechanisms

HIV has been previously reported to accelerate aging^33–35^. We were interested to discover whether this accelerated aging is due to cell-intrinsic or systemic effects. After validating our model on the Ren et al. (2021) COVID-19 dataset and exploring the effects of T cell changes with age and disease, we next turned to another external dataset comprised of patients infected with human immunodeficiency virus (HIV) on antiretroviral therapy and healthy controls^36^.Of these donors, 7 had HIV and were on antiretroviral therapy (ART), 1 had HIV but was off ART for the first visit, and 6 donors were healthy.

We applied our cell type prediction model to this dataset, successfully identifying and clustering the six distinct T cell types (Figure 4a) and finding the clusters to be in accordance with known cell markers (Figure 4b). Next, we used the age prediction models on this HIV dataset to predict the age of individual cells (Figure 4c) and donors (Figure 4d). We observed a low accuracy for age prediction comparing predicted cell age to donor age (R = .14, MAE = 13.6, p < 1*10^−16^) but a strong accuracy when cell age predictions were averaged per donor and compared to chronological age (R = 0.66, MAE = 12.6, p = .002). Unlike the COVID-19 dataset, we did not observe significant changes in cell type proportions after adjusting for multiple comparisons (Figures 4e-j). To assess differences in predicted age between HIV + ART donors compared to healthy controls, we performed an analysis of the age residuals for each cell type and identified significantly accelerated ages in HIV+ patients in the naive CD8 population (Figure 4k, p = .01).

**Figure 4.**
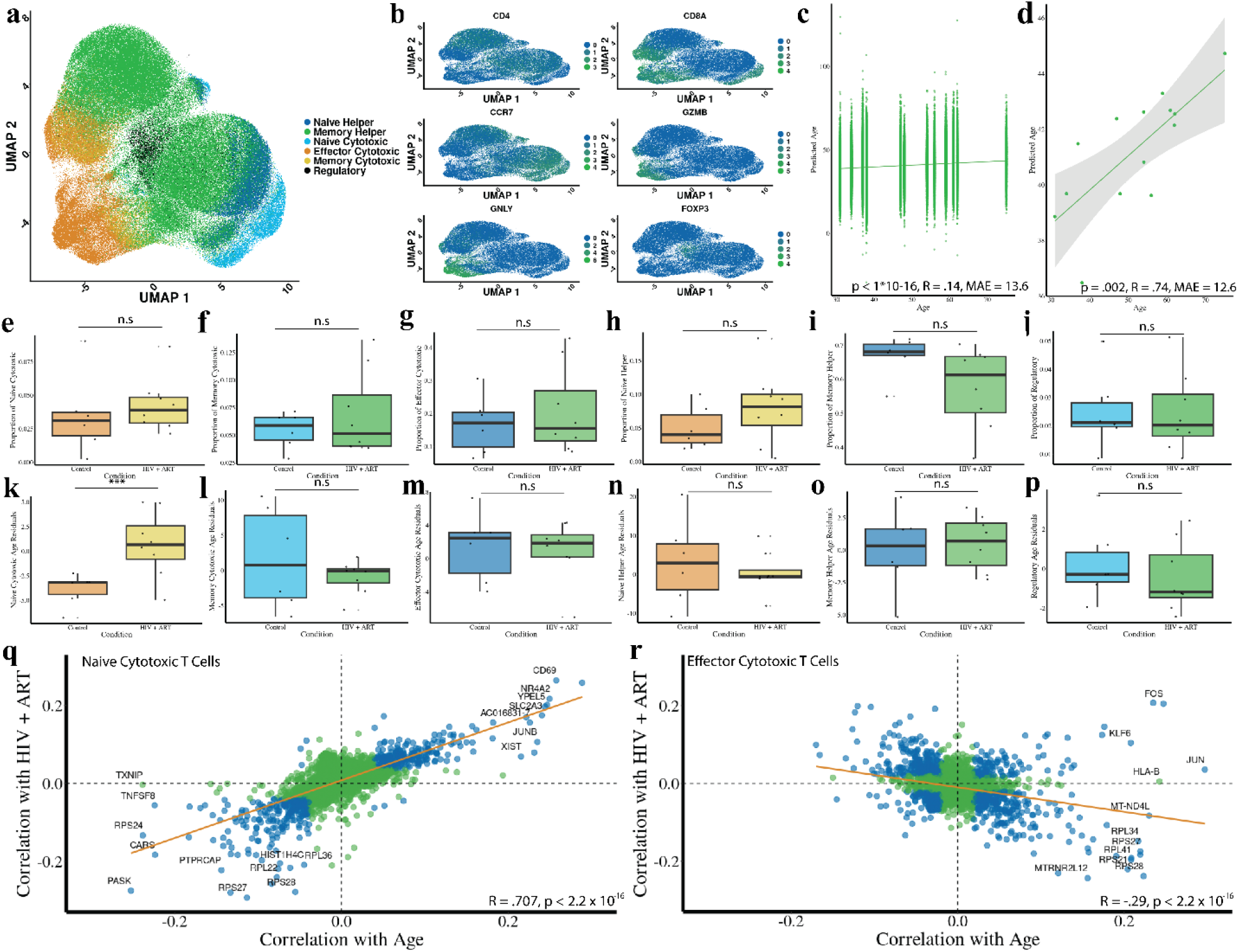
Impact of age and HIV + ART status on T cell composition, cellular aging, and gene expression. **a**, Predicted cell types using the cell type prediction model. **b**, Expression levels of canonical cell markers. **c**, Predicted age of individual cells versus donor chronological age. **d**, Mean prediction donor age across all cells compared to donor chronological age. **e-j**, Proportion of cells from COVID-19 donors and from healthy donors by cell type. **k-p**, Residual age prediction by condition for each cell type. **q**, Plot comparing genes correlated with HIV+ART and aging in naïve cytotoxic T cells. Blue represents a statistically significant effect. Orange line represents the mean regression. **r**, Plot comparing genes correlated with HIV+ART and aging in effector cytotoxic T cells. Blue represents a statistically significant effect. Orange line represents the mean regression. *** Bonferroni-corrected p-value less than or equal to .001, * Bonferroni-corrected P-value less than or equal to .05, # Bonferroni-corrected P-value less than or equal to .1.

Similarly to the COVID dataset, we performed an analysis of the transcriptional signature of HIV+ART compared to aging. In naïve CD8 cells (Figure 4q), we observed a strikingly strong correspondence between genes that had a partial correlation with aging and those that had a partial correlation with HIV+ART (R = .7, p < 1*10^−16^). Interestingly, many of the genes most strongly correlated with both HIV+ART and aging have been previously characterized to be a signature of HIV, such as CD69^37^. We also found a weaker negative correlation between genes associated with aging and HIV in CD8 effector cells (Figure 4r, R = -.29, p < 1*10^−16^).

### Biological insights into aging using the six T cell type-dependent model coefficients

One of the distinct advantages of using scRNA-Seq data or transcriptomics in comparison to other measurements of aging is the proximity to readily identifiable changes in biological function. In other clocks, such as those built using DNA methylation data, the interpretability of the clocks to gain information about the biology of aging is often limited^38^. We initially focused on the subset of genes used by all age predictions models to conduct a broad exploratory analysis. Accordingly, we performed a GO enrichment analysis for each individual T cell model, based on their relative weights in relation to the total gene set used in the model (Figure 5a). This allowed for an assessment of the significance of each gene within its respective model, providing a detailed understanding of the functional contributions of genes in different T cell types that are involved in age prediction. Both the naive cytotoxic T cells and the naive helper T cells have especially high involvement of ribosomal pathways in their biological processes analysis. We then investigated the 188 genes shared across all six models. GO enrichment using these 188 shared genes revealed cytosolic small ribosomal subunit, tumor necrosis factor receptor binding, small ribosomal subunit, and cytosolic ribosome as top hits across all cell type age prediction models (Figure 5b).

**Figure 5.**
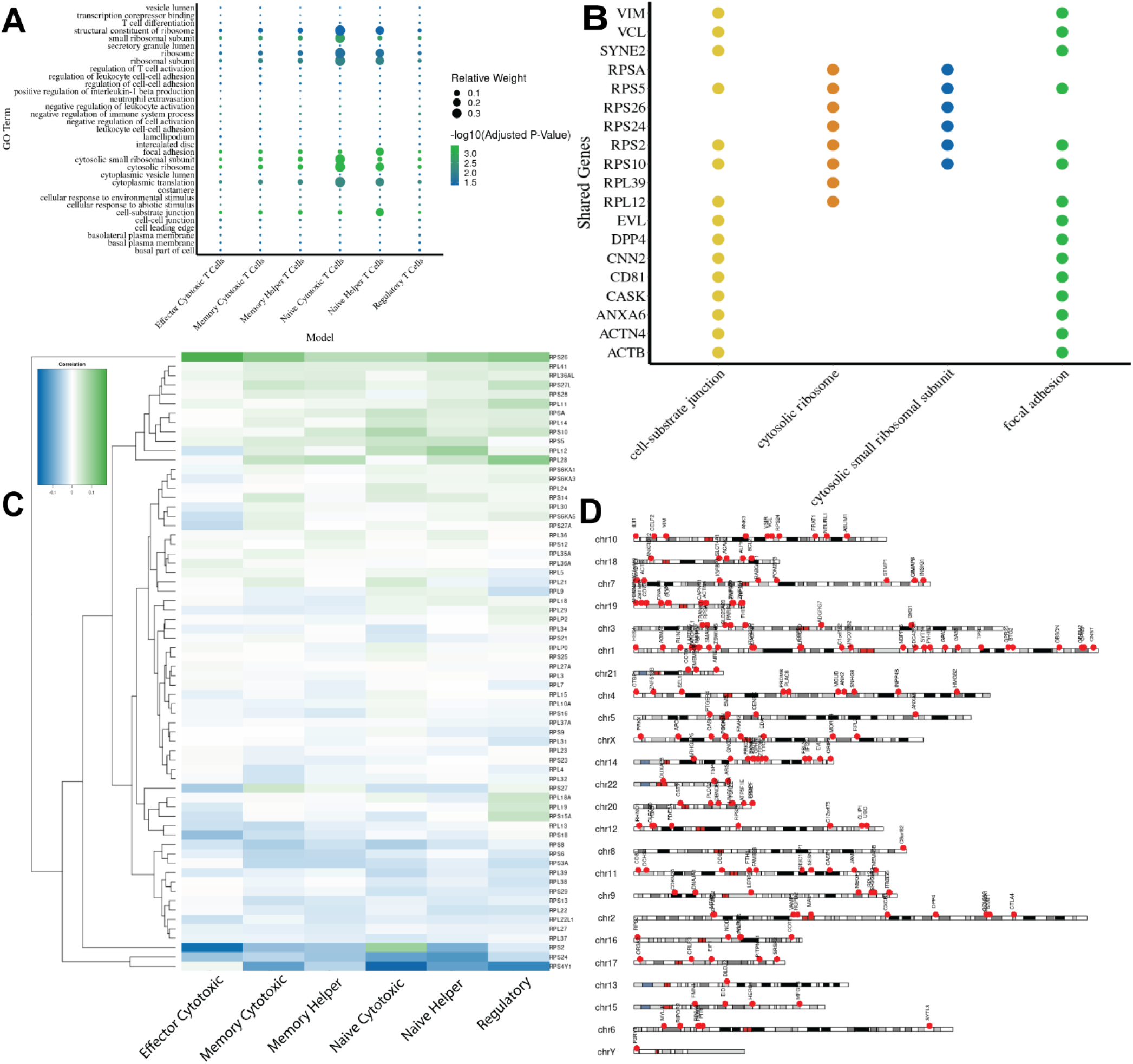
Drivers of single-cell transcriptomic clock age prediction. **a**, GO term enrichment investigating the shared genes across all six models. The black dotted line represents significance. Using a p-value of 0.05, the threshold for significance for the -log10 is approximately 1.3. **b**, A bar plot for the top four most enriched go terms based on the adjusted p-value. The top four genes from each GO term were then extracted and a comparison was made across all enrichment terms. **c**, Heatmap of correlations of each ribosomal gene with aging across each individual T cell subset. **d**, A karyoplot mapping all of the shared genes across all six models, making specific locations across human chromosomes.

Given the importance of ribosomal genes to the model, we were interested in investigating ribosomal gene expression with age across cell types (Figure 5c, Supplemental Figure 3). We found that most ribosomal genes have a similar pattern with age across all six cell types, although there are a few exceptions (such as RPS2). We were also interested in whether or not the predictive genes localized to particular regions of the genome. Karyoplot mapping of the specific chromosomal locations of these genes showed broad distribution across all chromosomes with a couple notably dense regions in chromosomes 1 and 14 (Figure 5d).

Due to increasing evidence for the role of defective translation of longer transcripts associated with aging^39,40^, we sought to identify whether such defects could be identified in the immune system during human aging using our single-cell transcriptional biomarker. We found that older individuals had, on average, shorter transcript lengths in every T cell subset except for naive CD4 T cells and memory CD8 T cells (Figures 6a-f). We were also interested in discovering whether shorter transcript lengths led to an accelerated predicted age. To test this possibility, we estimated transcript lengths in cells in the top and bottom deciles of predicted age relative to their chronological age. In memory helper cells and regulatory T cells, we found that donors with longer mean cellular transcript lengths were predicted to have a younger age relative to their chronological age (Figures 6g-6i). This is in agreement with previous reports linking mean cellular transcript length with aging^39,40^.

**Figure 6.**
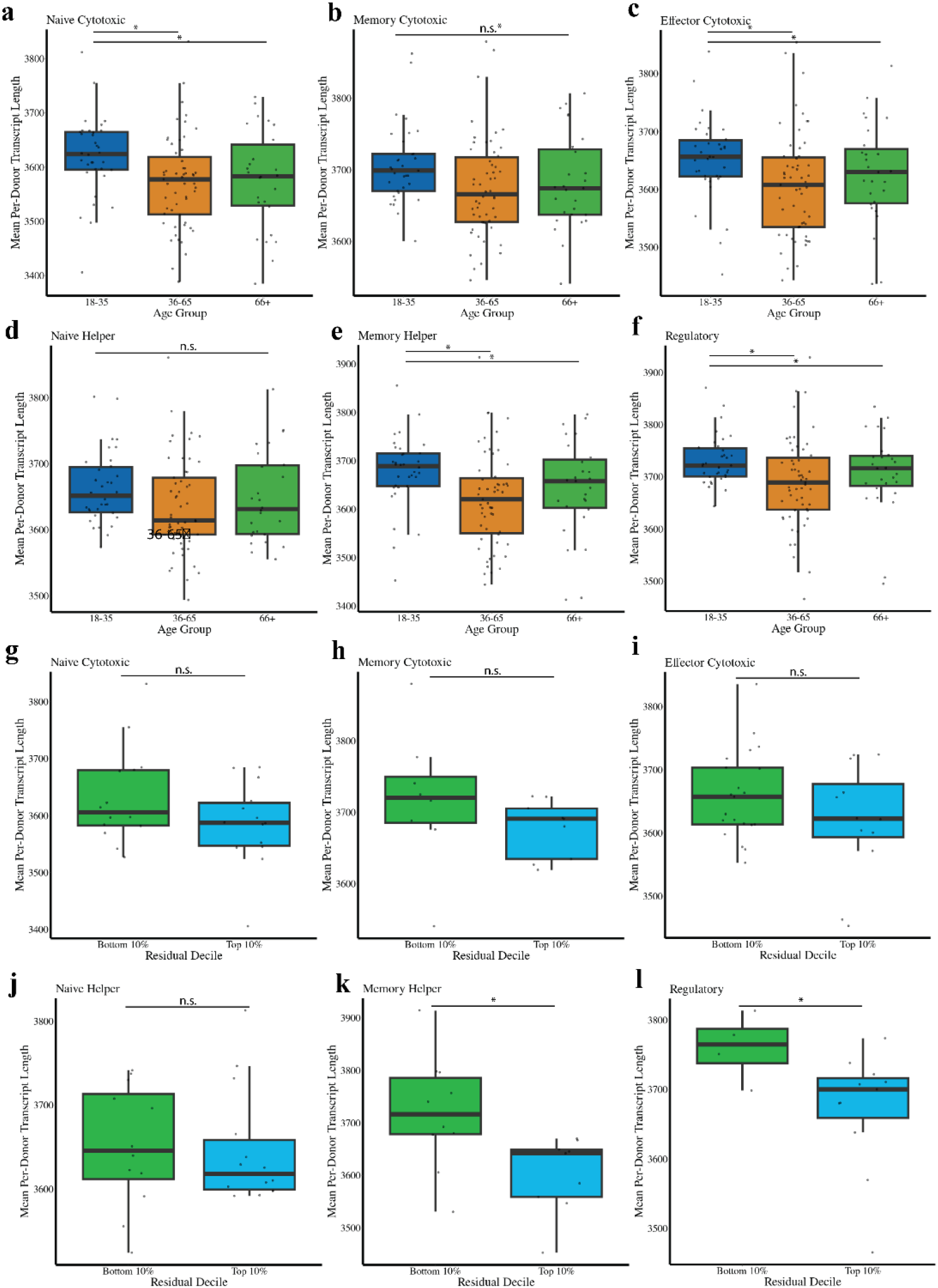
Mean transcript length is associated with aging in T cells. Mean donor transcript length per predicted cell type and age for **a**, naïve cytotoxic T cells, **b**, memory cytotoxic T cells, **c**, effector cytotoxic T cells, **d**, naïve helper T cells, **e**, memory helper T cells, and **f**, regulatory T cells. **g-l**, Comparison of the top 10% and bottom 10% deciles based on predicted age acceleration relative to chronological age in terms of mean transcript length for **g**, naïve cytotoxic T cells, **h**, memory cytotoxic T cells, **i**, effector cytotoxic T cells, **j**, naïve helper T cells, **k**, memory helper T cells, and **l**, regulatory T cells. *** Bonferroni-corrected p-value less than or equal to .001, * Bonferroni-corrected P-value less than or equal to .05, # Bonferroni-corrected P-value less than or equal to .1.

## Discussion

The immune system ages on a molecular, cellular, compartmental, and systemic level^41^. Molecularly, CD8 T cells lose chromatin accessibility at gene promoters, leading to reduced capacity for oxidative phosphorylation^42,43^. Cellularly, T cells progressively accumulate dysfunctional mitochondria^44^ and develop mis-calibrated receptor signaling pathways^45^. Thymic involution leads to a gradual reduction in naïve T cells^46^. On a systemic level, the immune system plays a key role in the senescence-associated secretory phenotype (SASP)^47^, and clonal expansion leads to an increased risk of leukemia^48^. Due to this complexity, it is critical to be able to study how aging affects multiple levels of immune organization simultaneously.

Furthermore, it is important to understand which aspect of immune aging a disease or intervention affects. Both acute COVID-19 infection^49^ and HIV^34^ have been reported to accelerate epigenetic aging, and antiretroviral therapy may slow it^35^. However, without a precise understanding of how each disease influences specific aspects of immune aging, it becomes difficult to develop targeted restorative therapies. Understanding the interplay between different levels of immune aging is also important for comprehending the basic biology of time-dependent T cell dysfunction.

By developing six age prediction models based on unique T cell-type dependent subsets, we can predict cell type and age of individual T cells using single cell transcriptomic data. We opted to automate cell type prediction in addition to age prediction, increasing the robustness and reproducibility of our findings. Our approach suggests that acute COVID-19 is both a cell-intrinsic and systemic immune aging phenomenon, while long-term HIV with ART treatment maintains an accelerated intrinsic aging signature on naïve CD8 T cells. We further showed that shorter mean transcript length is a feature of aging, and one that our transcriptomic clock tracks.

There are several additional interesting findings we discovered in this study. It is noteworthy that most of the genes used for age prediction are shared between specific cell type clocks (Figure 5A). This may be due to a core aging signature conserved between T cell subsets, though we are limited in our interpretation due to the sparseness of single-cell transcriptomics. It is also somewhat surprising that the T cell type most associated with accelerated aging in both acute COVID-19 and long-term HIV is naïve CD8s. Our findings are in accordance with previous literature showing a short-term loss of naïve CD8s in acute COVID-19^50^, but the current knowledge on cell-intrinsic dysfunction of naïve CD8s in either disease context is limited. Lastly, it is a unique finding that several ribosomal genes that are upregulated during aging are downregulated in the context of HIV + ART (Figure 4R). This aligns with previously known knowledge regarding HIV downregulating genes involved in ribosomal biogenesis^51^.

Though our age prediction models show significant promise, there are some limitations. Due to the sparse and noisy nature of single-cell RNA sequencing data, accurately predicting age to an accuracy similar to that of epigenetic clocks is challenging. Techniques such as Buckley et al.’s (2023) bootstrapping method reduce transcriptional variation and enhance cell prediction accuracy, though at the cost of losing single-cell resolution. Additionally, our models were validated using two specific external datasets from COVID-19 and HIV patients. While providing unique biological insights, they may not capture disease states and aging in a broader context. Further validation using more varied and diverse datasets would prove beneficial and ensure broader generalizability of our findings. Lastly, although we identify and characterize cellular ribosomal signatures of aging in a unique manner using our novel biomarker, it is likely that there is technical enrichment of a ribosomal signature in scRNA -Seq aging datasets due to the general high expression of ribosomal genes.

During preparation of this manuscript, multiple reports were released complementing findings discovered in this study. Another group developed a single-cell transcriptomic clock for human PBMCs, though interestingly they found that COVID infection had a *rejuvenative* effect on T cells^52^. A separate work similarly identified an opposite aging effect based on single-cell vs. bulk transcriptomic readouts^53^, reinforcing the importance of understanding the effect of cell type composition on aging measurements.

In summary, our study provides a novel framework for understanding immune aging by integrating single-cell transcriptomic data with automated T cell type and age prediction. By highlighting both cell-intrinsic and systemic aspects of immune aging, our approach offers insights into how diseases like COVID-19 and HIV differentially impact the immune system at the cellular level. These findings highlight the potential of single-cell aging biomarkers to improve the specificity of aging diagnostics and to inform therapeutic strategies aimed at mitigating age-related immune decline. Future research should focus on expanding this model to other cell types and disease conditions to build a more comprehensive map of aging across the immune system, and to explore how therapeutic interventions could potentially reverse or modulate these aging signatures. Ultimately, our work is a contribution to understanding how aging and disease impacts the immune system across different layers of organization.

## Methods

### Human cohorts

The Terekhova et al. (2023) dataset was used to train the cell type prediction and age prediction models. This dataset consists of roughly 2 million peripheral blood mononuclear cells from 317 samples from 166 individuals aged 25-85 years old. All participants were healthy Caucasian non-obese non-smokers. Blood was collected between 2018 and 2021 after an overnight fast. The Chromium Single Cell 5’ v2 Reagent Kit from 10x Genomics was used to generate single-cell transcripts. We utilized the Yasumizu et al. (2024) dataset to validate the accuracy of our cell type prediction and age prediction models. This dataset consists of 1.8 million CD4+ T cells generated using the Chromium Next GEM Single Cell 5′ Kit v2.

The Ren et al. (2021) dataset made up of 284 samples from 171 COVID-19 patients and 25 healthy individuals was used as a validation data for our model. The samples from COVID-19 patients were categorized into moderate convalescence (n = 89), moderate progression (n = 33), severe convalescence (n = 51) and severe progression (n = 83) according to severity and stage. They then generated a transcriptomic dataset of 1.46 million immune cells using Chromium Single Cell 3’ v2 Reagent, Chromium Single Cell 3’ v3 Reagent, Chromium Single Cell 5’ v2 Reagent, and Chromium Single Cell V(D)J Reagent kits from 10x Genomics. For our analysis, we included only cells derived from patients who had COVID-19 within two weeks of sample collection. Furthermore, we limited our analysis to cells processed with the Chromium Single Cell 5’ V2 kit.

The Wang et al. (2024) dataset consists of fourteen donors, eight of whom have HIV. Of the eight individuals with HIV, seven are actively treated with ART. The dataset consists of 262,818 PBMCs from these donors generated using the Chromium 5′ Single Cell Gene Expression system from 10X Genomics.

### Data processing

All datasets were filtered to remove cells with high mitochondrial reads and low transcript counts (< 2000 reads). Datasets were then processed using the Seurat^54^ pipeline, using the NormalizeData, ScaleData, FindVariableFeatures, RunPCA, FindNeighbors, FindClusters, and RunUMAP functions to identify cell types. For datasets including other peripheral blood mononuclear cell subpopulations, cells were filtered to include only T cells. The original Terekhova et al. (2023) dataset used for clock training was filtered to include only genes that had at least a > 0.01 or a < -0.01 correlation with age. All other datasets were filtered to include only genes used for cell type and age prediction.

### Cell type annotation

Cell type annotation was conducted using the ‘Seurat’ package (v5.1.0) in R. Gene expression matrices were loaded and aligned with the metadata. T cell subsets were re-annotated for more specificity and clustering into six T cell subsets. Cell clustering was performed with the ‘FindNeighbors’ and ‘FindClusters’ functions from Seurat (34) followed by visualization using a UMAP. Clusters were manually annotated and renamed based on marker gene (CD8A, CD4, CCR7, GZMB, GNLY, FOXP3) expression. The re-labeled T cell subsets were incorporated into the metadata and used for training the cell type prediction model.

### Cell type and age prediction models

The dataset was split into training and test sets based on donor ID. The split was 80-20 with 80% of the donor IDs being saved in the training dataset and 20% being saved in the test dataset. For cell type prediction, a multinomial logistic regression model was trained using the ‘glmnet’ package in R. Six models were then built for age prediction—one for each T cell cluster—using the Terekhova et al. (2023) data. Each model then performed elastic net regression using the ‘glmnet’ package with the cv.glmnet function and the alpha parameter set to 0.5. Non-zero coefficients from the elastic net model were extracted and then used to fit a second model, as previously described^55^.

### Enrichment analyses

Gene ontology (GO) term enrichment analysis was performed on shared genes using the ‘enrichGO’ function from the ‘clusterProfiler’ package in R^56^. A chromosome visualization plot was created using the ‘karyoploteR’ package in R where the gene locations were plotted on a karyoplot^57^. The genes plotted were genes used as coefficients across all six T cell models.

### Statistics

To determine the statistical significance of the differences in residuals between conditions, pairwise t-tests were performed with Bonferroni correction to adjust for number of cell types analyzed. ANCOVA tests were utilized to measure the differences in cell proportions while accounting for the confounding effect of age by adjusting for age-related variability. The ‘aov’ function was used from the ‘stats’ package in R. Additionally, the ‘emmeans’ package was used to provide adjusted mean estimates and p-values to compare between conditions^58^. The ‘ggplot2’ package was utilized for graphing.

For gene expression, residual, and cell proportion analyses, cells were sampled to maintain the same number of cells per donor. We performed ANCOVA to compare proportions between conditions (Control vs. HIV) while adjusting for age. Statistical analysis was performed using ‘emmeans’^58^ for pairwise comparisons and car for variance inflation factor (VIF) calculations. Partial correlations were calculated to control for confounding factors (donor age or condition) for gene expression analysis using the ‘ppcor’ package.

## Data Availability

The Terekhova et al. (2023) data is available through the Synapse platform (syn49637038). The Ren et al. (2021) data can be found on the Gene Expression Omnibus (GSE158055). The Wang et al. (2024) dataset is accessible from the Gene Expression Omnibus (GSE243905).

## Code Availability

The code used to build the model and perform all analyses will be publicly available on GitHub prior to publication. The primary programming languages used were R and Python with freely available software packages. All computational work was performed on Amazon Web Services (AWS) Elastic Compute Cloud (EC2) instances.

## Acknowledgements

This project was supported by institutional funding from the Buck Institute for Research on Aging.

## Author Contributions

A.T., and S.L. jointly conducted the work described in this study, including drafting and writing the manuscript. All authors contributed to manuscript editing and revising.

## Competing Interests

There are no competing interests to declare.

## Notes

### Competing Interest Statement

The authors have declared no competing interest.

