## Supplemental Figures for "Utilizing blood single-cell transcriptomics to integrate intrinsic and systemic immune aging"

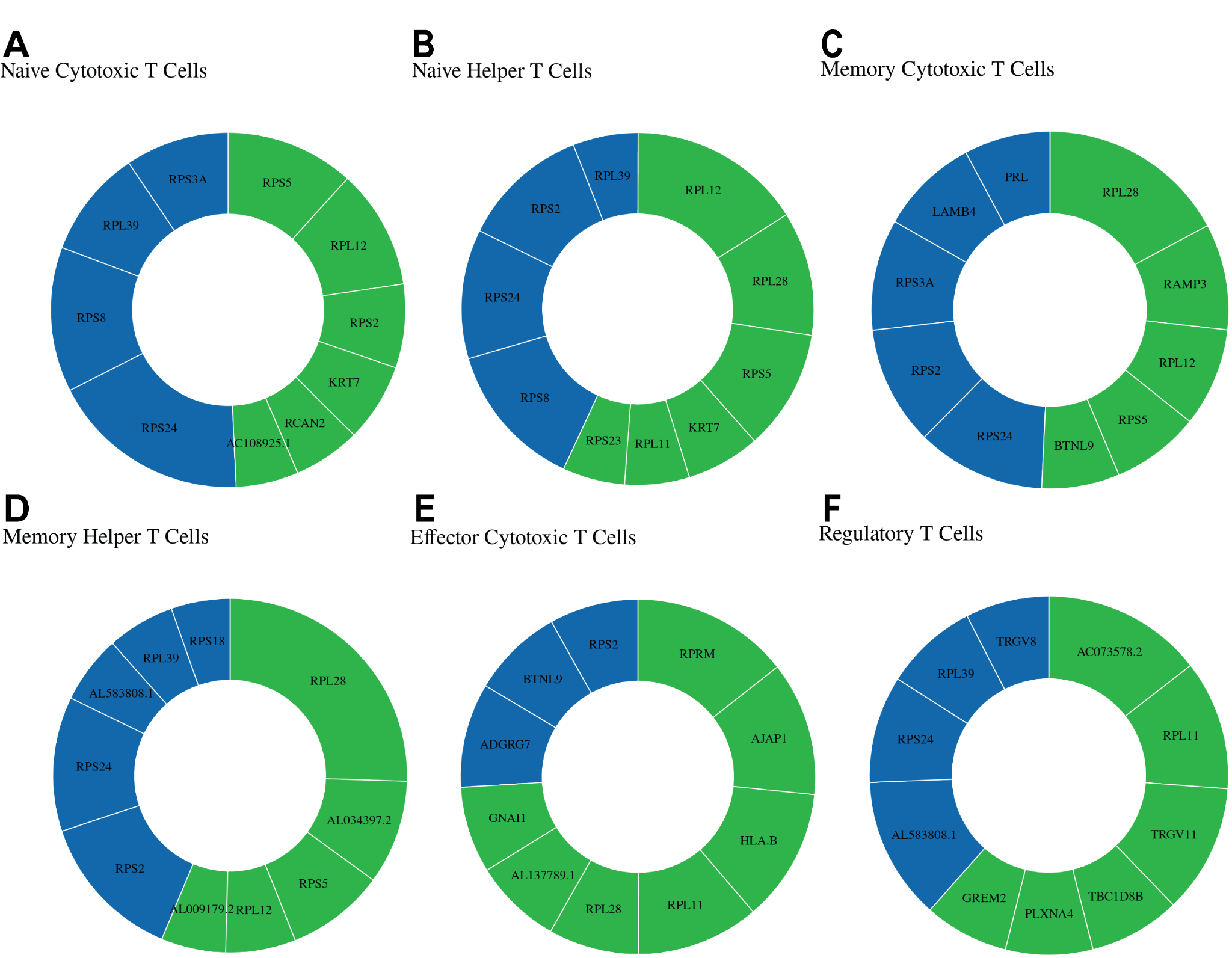

**Supplemental Figure 1. Significant gene contributions to each T cell type-dependent model.** Donut plots for each model using the top ten most significant coefficients per T cell type-dependent model. Green represents upregulated genes and blue represents downregulated genes.

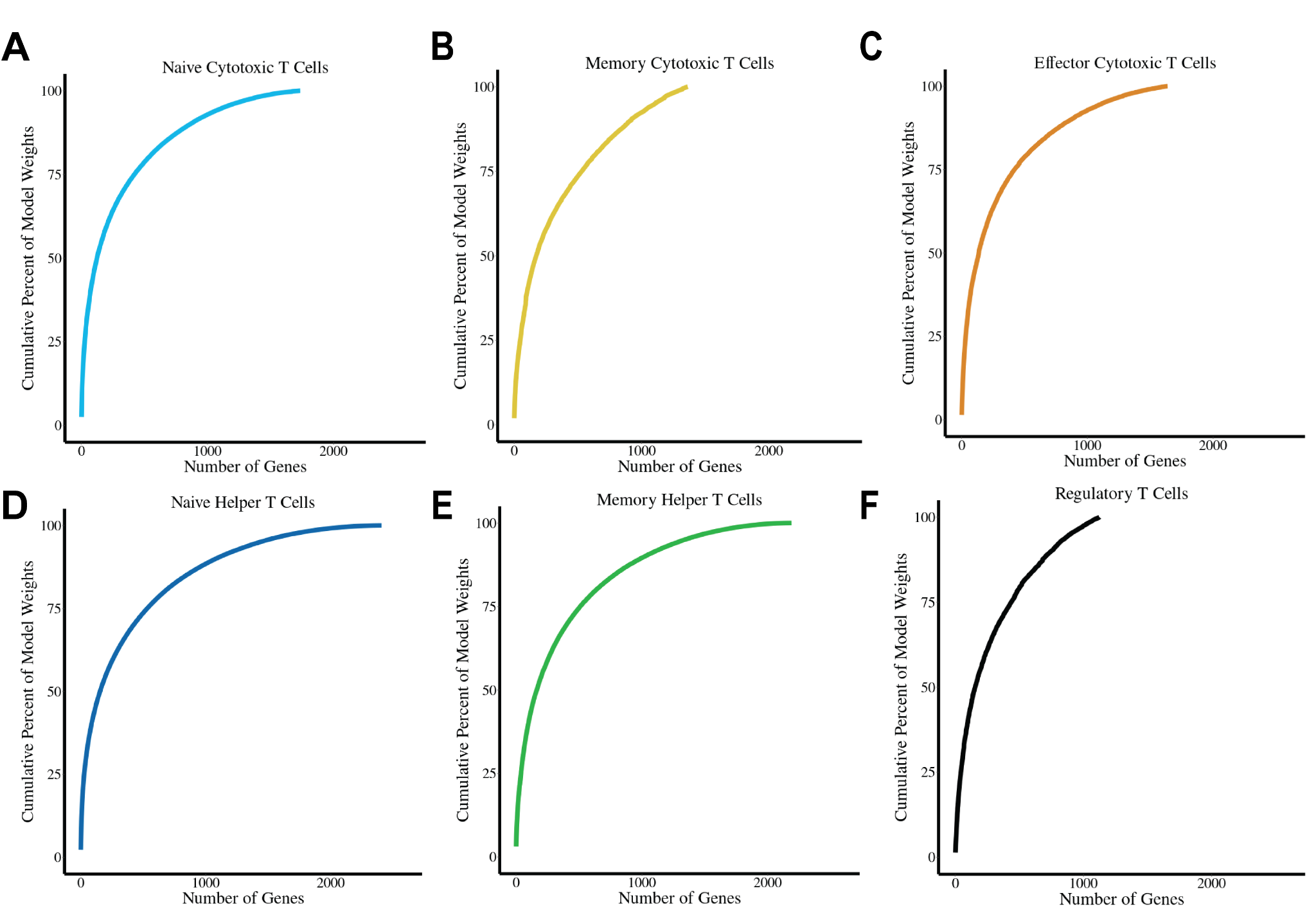

**Supplemental Figure 2. Cumulative percent of model weights.** Cumulative distribution plots of model weights assigned to genes for each T cell subset. Each panel shows how the cumulative percentage of the total model weights is distributed across all of the genes used in the model as a cumulative distribution function.

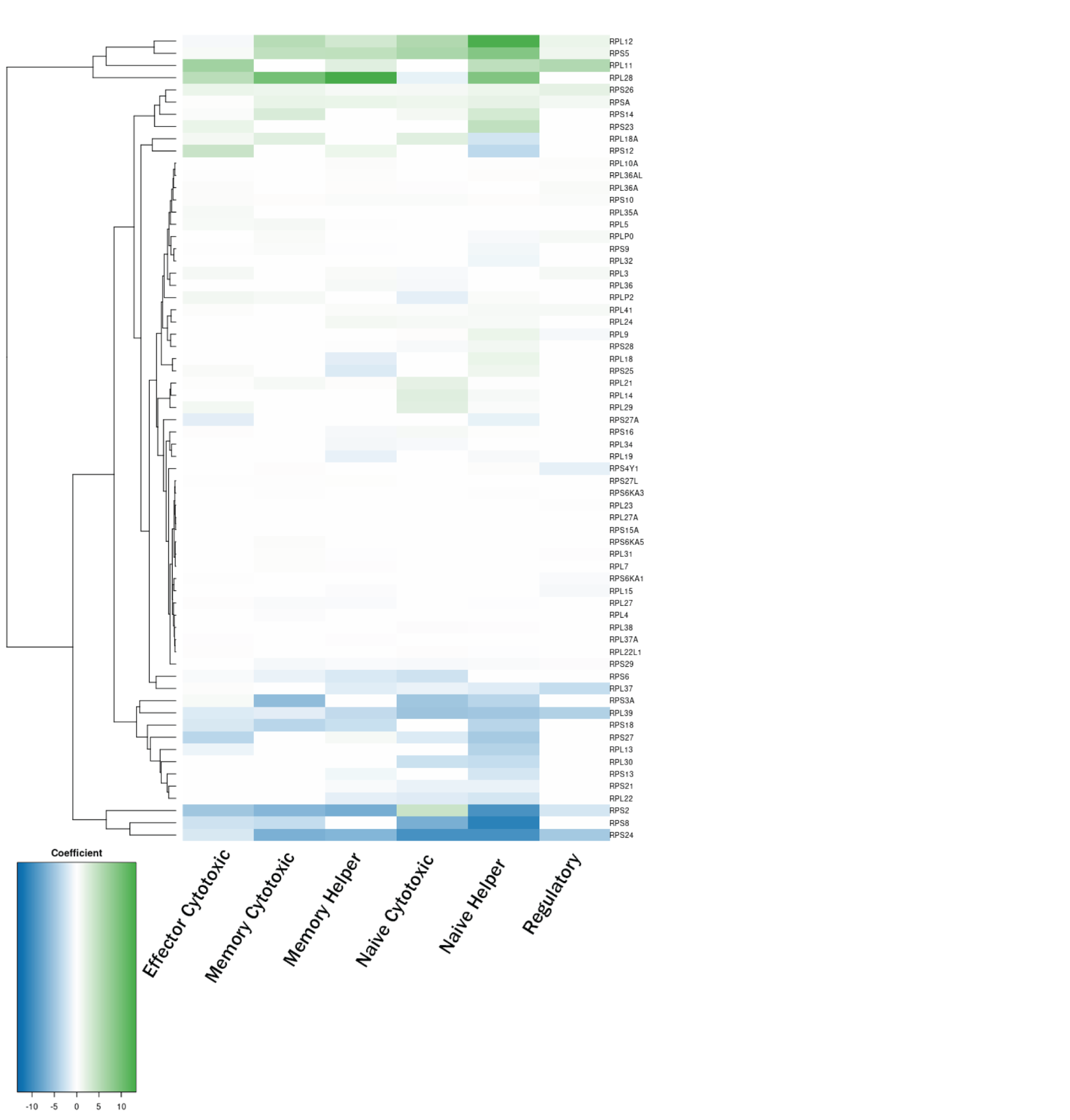

**Supplemental Figure 3. Ribosomal coefficient mapping across cell types.** A heatmap for the Terekhova et al. (2023) dataset, with the age prediction coefficients across ribosomal genes for every cell type.
